# Auditory cues modulate the short timescale dynamics of STN activity during stepping in Parkinson’s disease

**DOI:** 10.1101/2023.10.31.565053

**Authors:** Chien-Hung Yeh, Yifan Xu, Wenbin Shi, James J. FitzGerald, Alexander L. Green, Petra Fischer, Huiling Tan, Ashwini Oswal

## Abstract

**Background:** Gait impairment has a major impact on motor performance and quality of life in patients with Parkinson’s disease (PD). The neurophysiological mechanisms of gait impairment remain poorly understood, meaning that treatment options are currently limited. It is believed that basal ganglia oscillatory activity at β frequencies (15-30 Hz) may be a contributor to gait impairment in PD, but the precise dynamics of this oscillatory activity during gait remain unclear. Auditory cues are known to lead to improvements in gait kinematics in PD. If the neurophysiological mechanisms of this cueing effect were better understood they could be leveraged to treat gait impairments using brain-computer interface (BCI) technologies.

**Objective:** We aimed to characterize the dynamics of subthalamic nucleus (STN) oscillatory activity during stepping movements in PD and to establish the neurophysiological mechanisms by which auditory cues modulate gait.

**Methods:** We used local field potentials (LFPs) to study STN oscillatory activity in eight PD patients while stepping in place with auditory cueing. Hidden Markov Models (HMMs) were used to discover dynamic brain states that occurred pre-sound, on-sound, and post-sound cues.

**Results:** The occurrence of β bursts was suppressed during and after auditory cues. This manifested as a decrease in their fractional occupancy and state lifetimes. Interestingly, α transients showed the opposite effect, with fractional occupancy and state lifetimes increasing during and after auditory cues.

**Conclusions:** We found transient oscillatory states in the STN LFP during stepping and showed that α and β oscillations are differentially modulated by auditory cues during stepping in PD.

## 1. Introduction

Gait impairment is a major cause of disability in Parkinson’s disease (PD)^1^. Gait disturbances confer an increased risk of falls, in turn potentially leading to morbidity related to injury and a loss of both independence and quality of life^2,3^. Traditional therapeutic approaches for PD, including dopaminergic drug therapy (e.g., levodopa) and deep brain stimulation (DBS) of basal ganglia structures (e.g., subthalamic nucleus (STN) or globus pallidus), are often insufficiently effective for treating gait symptoms and may even lead to a worsening of gait^4-6^. There is consequently a need to develop a better understanding of the neurophysiology of gait impairments, so that improved therapies can be developed.

Dynamic oscillatory activity within complex brain structures is believed to reflect both physiological and pathophysiological processes^7^. Recent studies of patients undergoing DBS surgery have revealed how synchronized oscillatory activity across the cortico-basal ganglia circuit can contribute to gait kinematics and motoric impairments in PD^8-11^. For example, it has been shown that excessive STN activity at beta frequencies (15-30 Hz), which is traditionally thought to be related to both bradykinesia and rigidity, can also precede paroxysmal episodes of freezing of gait (FOG). Consistent with this observation is the finding that both motor cortical and STN beta activity display gait phase-locked modulations during walking^9^. Interestingly, cortico-STN synchronisation at lower frequencies within the alpha band (8-12 Hz) is also observed during gait execution, with FOG episodes being reportedly accompanied by transient decoupling of alpha synchronisation^11^. These findings suggest opposing roles for alpha and beta synchronisation in gait control.

One way of improving gait coordination in PD is to present rhythmic visual or auditory cues during stepping^12-15^. Auditory cueing may enhance both walking speed and stride length, in addition to ameliorating FOG episodes ^14-23^. Although we have previously shown that cueing to the timing of a heel strike can modulate STN beta activity, it remains unclear whether cues can modulate STN activity at shorter timescales^9^.

Establishing the dynamics of STN activity at short timescales during stepping could be important for identifying biomarkers of both normal and abnormal gait in PD. The discovery of oscillatory signatures that reliably reflect clinical state is of great interest, given that these signals could be selectively modulated (either amplified or suppressed) using adaptive DBS (aDBS) regimes^24,25^.

Traditional approaches for identifying spectral activations at short timescales (known as *bursts*) rely on the prior specification of an amplitude threshold, which defines the onset and offset of a burst^26-28^. This method becomes challenging when attempting to define bursts over multiple participants and frequency bands^29,30^. In this paper, we overcome these limitations by using an unsupervised machine learning approach, Hidden Markov Modelling (HMM)^31^, to detect bursting dynamics within the STN during stepping in patients with PD. We hypothesized that: (1) STN bursting activity within the alpha and beta bands would be differentially modulated during stepping, in keeping with the notion that activity within these frequency bands has opposing effects on gait, and (2) that auditory cues enhance gait-related modulation of activity within these two frequency bands.

## 2. Materials and Methods

### 2.1 Patients and Experiments

Eight PD patients (7 males and 1 female) with bilateral STN DBS electrodes were included in this study (clinical details in Table 1). These data have been used to study the effects of rhythmic auditory cues (sound on and sound off) on STN β modulation during periodic alternating stepping in place while sitting, and behavioral analyses (step-to-cue difference and step-to-interval duration) were performed to measure the synchronization of stepping with auditory cues^9^. In this study, we further subdivided the auditory conditions (pre-, on, and post-sound) and explored the short timescale dynamics of STN activity in multiple bands.

**Table 1.** Clinical information of 8 subjects with PD having periods in sound stimulation, including age, gender, main symptoms, disease duration, UPDRS-III score, and DBS lead.

| SUB ID | Age | Gender | Main Symptom | Disease Duration (yrs) | UPDRS-III OFF/ON Levodopa | Levodopa equivalent dose (mg/d) | DBS Lead |
| --- | --- | --- | --- | --- | --- | --- | --- |
| 1 | 59 | M | tremor, bradykinesia, dyskinesia | 7 | 53/18 | 1195 | Boston Scientific DB-2202 |
| 2 | 64 | F | rigidity, tremor, FOG | 16 | 66/36 | 1628 | Boston Scientific DB-2202 |
| 3 | 59 | M | fluctuations, tremor | 14 | 36/8 | 1062 | Medtronic3389 |
| 4 | 56 | M | fluctuations, dyskinesia | 7 | 42/26 | 1365 | Medtronic3389 |
| 5 | 62 | M | tremor, rigidity, dyskinesia, mild FOG | 12 | 59/15 | 1000 | Medtronic3389 |
| 6 | 71 | M | tremor, FOG | 15 | 36/18 | 785 | Boston Scientific DB-2202 |
| 7 | 61 | M | rigidity | 9 | 33/11 | 1293 | Medtronic3389 |
| 8 | 59 | M | tremor, mild FOG | 10 | 28/8 | 1010 | Boston Scientific DB-2202 |

M, male; F, female; FOG, freezing of gait.

Patients had an average age of 61.4±4.6 yrs (range 56– 71 yrs) and an average disease duration of 11.3±3.5 yrs (range 7–16 years). The mean Unified Parkinson’s Disease Rating Scale (UPDRS) scores (part III) were 44.1±13.6 (range 28–66) and 17.5±9.6 (range 8–36) in the off-levodopa and on-levodopa states. All experimental procedures were approved by the local ethics committee with informed written consent being sought from all participants. LFP recordings were performed in the on-levodopa state, 3-7 days after electrode implantation surgery.

During recordings, patients were asked to sit in a comfortable chair with their arms resting on their laps. Two flat-plate pressure sensors were placed on the floor to record right- and left-foot stepping movements. An instructional walking video that looped after each stepping cycle (alternate left and right foot heel strike, separated by a delay of 1 second) was displayed on a laptop monitor in front of the patient. Patients were instructed to synchronize the timing of their footsteps to match the timing of the footsteps of the video character. In addition to the visual cue, a metronome sound was provided at the time of each heel strike displayed in the video. This provided additional information about the timing of heel strikes via the auditory system. The recorded LFP could then be distinguished into pre-sound, on-sound, and post-sound conditions. LFP recordings were performed during stepping while sitting to avoid movement artefacts.

A TMSi Porti amplifier (TMS International) with a common averaged reference was used to record STN-LFPs in a monopolar configuration with a sampling rate of 2048 Hz. The data were re-referenced offline to obtain more spatially focal bipolar signals by subtracting the data from neighboring electrode contacts. Prior to further analysis, recorded signals were downsampled to a frequency of 1000 Hz.

### 2.2 Data Processing and Time-frequency Analysis

All data processing and analysis were performed in MATLAB (v2019a, MathWorks, Natick, MA). For each subject, the single bipolar STN LFP channel with the greatest mean power over the uniform distributed 8-35 Hz frequency range (including alpha and beta bands) was selected for further analysis. Raw LFPs were notch filtered at 50 Hz to suppress power line noise, and then high pass filtered at 8 Hz, using a sixth-order Butterworth filter. Data fragments with poor filtering effects were discarded by visual inspection (i.e., strong instrument noise or movement artifacts). Finally, signals were low pass filtered at 48 Hz (using a sixth-order Butterworth filter) before being downsampled to a frequency of 100 Hz. To observe how the frequency dynamics of a signal vary over time, the spectrogram of the preprocessed LFP was constructed by continuous Morlet wavelet transform with the wavelet cycle set to span six cycles^32^. The wavelet function is given by:

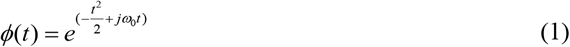

Then the wavelet function is scaled and shifted according to:

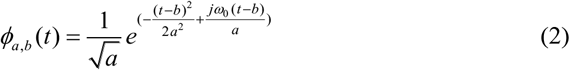

where *a* and *b* represent a scale factor and a time shift factor, respectively. The factor *a* controls the expansion and contraction of the wavelet function, larger values of a correspond to a higher frequency resolution. The factor b implements the shift of the wavelet function along the time axis.

### 2.3 Time-delay Embedded Hidden Markov Model

The Hidden Markov model (HMM) is an unsupervised machine learning approach for detecting transient states within the data such as spectral events or bursts^27,29,33-35^. The HMM assumes that data are generated from a hidden sequence of a finite number of states. At each time point, only one state is active and the data observed in each state are drawn from a probabilistic observation model, i.e., a probability distribution such as multivariate Gaussian. The HMM is a doubly stochastic process, meaning that noise is incorporated in both the state transitions and the observation model.

Important assumptions of the HMM are that: 1) the current hidden state depends only on the previous hidden state, and is unrelated to other time states and observations, and 2) the observed data at any time point depends only on the current hidden state and is unrelated to other states.

The HMM has three parameters, *π, A*, and *B* which are learned from the data. The occurrence probability of each initial state is denoted *π* = (*π*_1_,*π* _2_,…,*π* _*N*_), where the subscript *N* denotes the total number of states. The state transition matrix is given by *A*, where each entry *A*_*ij*_ corresponds to the probability of state *i* transitioning to state *j*. The matrix *B* represents the parameters of the observation model – which in the case of a multivariate Gaussian would include the mean and covariance for each state.

We used a specific variant of the HMM known as the time-delay embedded hidden Markov model (TDE-HMM). In this approach, an input signal is shifted by different time lags, *N* time lags refer to having *N* samples of either side to the current timepoint (i.e., a window of length in 2*N* +1), wherein the shifted signals are concurrently input as data in 2*N* +1 channels for HMM inference^27,35^. The purpose of this is to allow the model to infer states as periods of distinct autocovariance. This translates to transient episodes of distinct spectral content, known as bursts in the frequency domain^29^. In contrast to traditional threshold-based approaches, HMMs allow for the automatic detection of bursting activity across multiple frequency bands^26,27^.

TDE-HMM inference requires the number of states and time lags under consideration to be predefined. The number of time lags determines the window duration so a lag of means that the window includes data points on either side of that representing the time under analysis. Short windows allow for better temporal resolution at the expense of frequency resolution. Here, we performed inference for 12 states and 3 time lags. The TDE-HMM analysis was performed using the HMM-MAR MATLAB toolbox (https://github.com/OHBA-analysis/HMM-MAR).

### 2.4 Masking Empirical Mode Decomposition

Empirical mode decomposition (EMD) - known for its advantages for analyzing nonlinear and non-stationary signals - decomposes dynamical signals into a finite number of intrinsic mode functions (IMFs), wherein each IMF contains local characteristic information of the original signal at different time scales^36^. The input signal can therefore be expressed as the sum of multiple IMFs and a residual term:

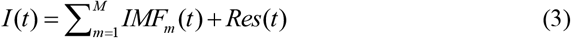

where *I* (*t*) is the input signal, *IMF*_*m*_ (*t*) denotes the *m*^*th*^ IMF component, and *Res*(*t*) stands for the Residual term.

Developments in EMD approaches include the masking EMD (MEMD) algorithm, which can overcome the issues of mode mixing and mode splitting^37-40^. More specifically, MEMD involves applying a sinusoidal masking signal. Following this, a desired frequency for each band is selected to improve MEMD^41,42,^ thereby allowing each IMF to capture the dynamics of a distinct frequency band of interest. In this study, the frequency bands of interest were the alpha (α, 8-12 Hz), low beta (Lβ, 13-21 Hz), high beta (Hβ, 22-35 Hz), and gamma (γ ∼ 40Hz) bands.

### 2.5 Assigning HMM States to IMF Frequency Bands

We used IMFs corresponding to the four frequency bands of interest, to select states identified from the HMM for further analysis. We correlated each state time course identified from the HMM with Hilbert envelopes of the α, low-β, high-β, and γ frequency activities obtained by MEMD (i.e., IMFs). HMM states with exclusively negative correlation coefficients were assigned to background activity (BG). States with positive correlations were assigned to the IMF frequency band with which they were maximally correlated. This procedure meant that more than one HMM state could be assigned to a single frequency band^27^.

### 2.6 Feature Extraction of HMM States

The above procedure was applied for the selected channel of all subjects. Spectral characteristics of identified state time courses were determined using the multitaper method^43^, with a taper smoothing frequency of 3.125 Hz (the time-half-bandwidth product was 4 and the total number of tapers used was 7).

Time domain features^44^ were also extracted from the TDE-HMM model for comparison across the three sound conditions. These included the fractional occupancy (FO) of each state, which was defined as the proportion of time that each HMM state occupied over the entire time sequence. The FO of a specific state *k* is given by:

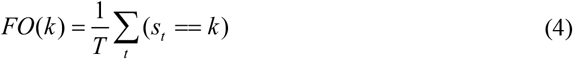

where *T* is the total number of time points, and *s*_*t*_ is the number of time points at which state *k* was active. State lifetime (LT) was defined as the average duration of each state, before transition to another state. For a specific state *k* this is defined as:

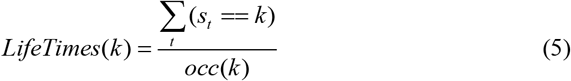

where *occ*(*k*) represents the number of occurrences of state *k*.

The transition probability (TP) from each state to every other state (including itself) was also obtained from the TDE-HMM model.

### 2.7 Statistics Analysis

Statistical analysis was performed in MATLAB (v2019a, MathWorks, Natick, MA). For comparing state features across frequency bands and sound conditions, we used the Wilcoxon signed-rank test, with α = 0.05. Multiple comparison correction was performed using the Benjamini-Hochberg false discovery rate (FDR) method on the comparisons between the three sound conditions in each frequency band. FDR-correction was also applied for the 25 comparisons of the transition probabilities. All results are presented in the form of box-and-whisker plots with a box from the first quartile to the third quartile and a horizontal line in the middle representing the median. The top and bottom whiskers link the maximum and minimum values.

## 3. Results

### 3.1 Decoding HMM States

We observed that the TDE-HMM effectively identifies transient spectral states in the STN LFP. Fig. 2 shows the results of state estimation for a 4-s long LFP in a single participant (Subject 4) during the experimental paradigm. The second panel in this figure displays the time course for each different state. For example, between 2.52-2.64 s, there was a spectral peak at ∼20 Hz which corresponded to a single state shown in the yellow line (black botted box). Between 2.95-3.04 s, there was a spectral peak at ∼36 Hz which corresponded to a single state shown in the purple line (black dotted box).

**Fig. 1.**
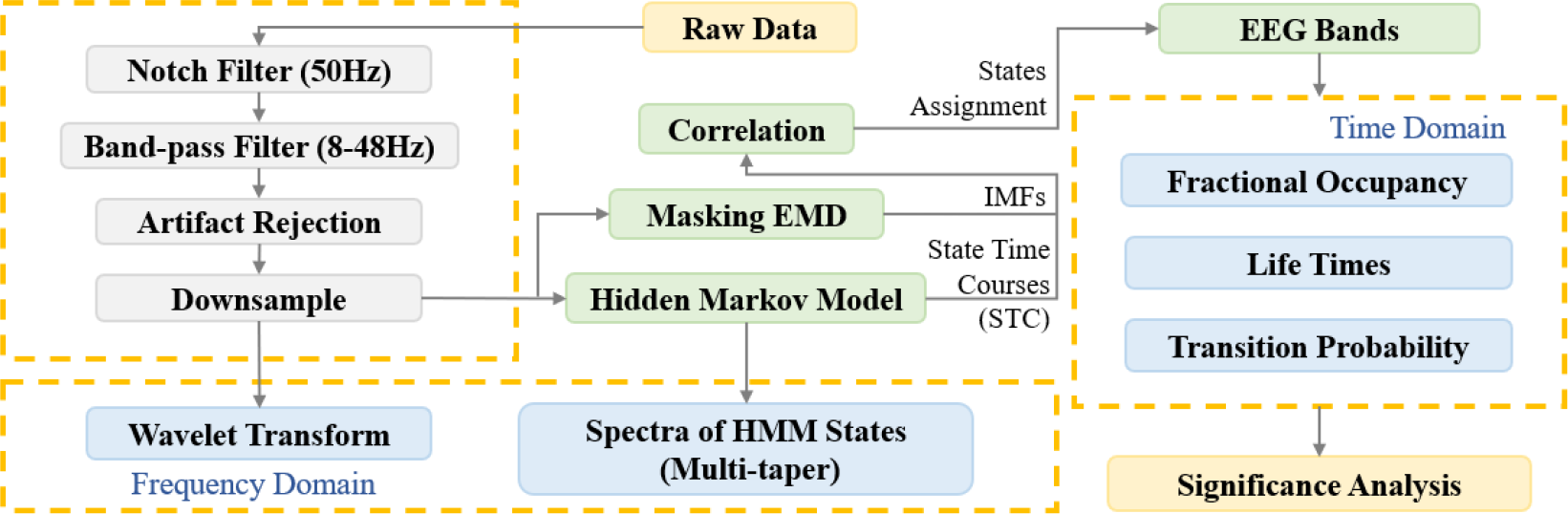
The algorithm flowchart of the LFP-based brain-state decoder.

**Fig. 2.**
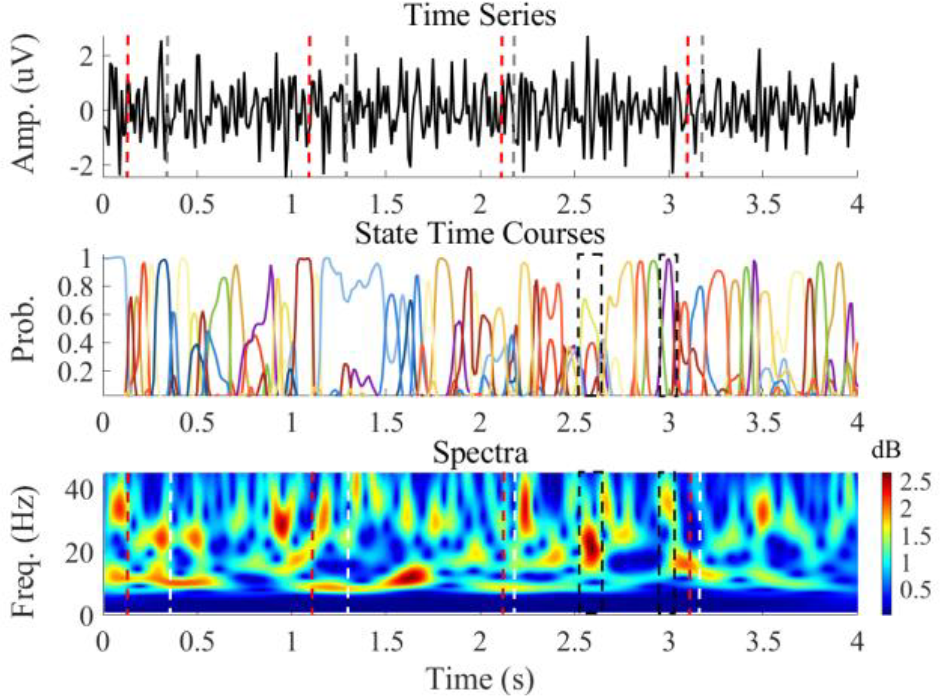
A 4-s long preprocessed LFP recording (Subject 4) which contains 4 steps. The top panel is the preprocessed LFP. The red dashed lines represent auditory cues, whilst the grey lines correspond to the stepping movements. The middle panel displays the probability time courses for the HMM states. The bottom panel shows the corresponding time-frequency spectra calculated using the wavelet transform. In this, the red dashed line represents auditory cues, whilst the white dashed line represents stepping movements.

### 3.2 Decomposition Performances using Masking EMD

Masking EMD was used to extract IMFs corresponding to the classical LFP frequency bands. Illustrative results of this decomposition for Subject 4 are shown in Fig. 3. The left-hand side of the figure shows IMFs corresponding to the canonical LFP bands (α, low-β, high-β, and γ), whilst the right-hand side displays the corresponding frequency spectra of the IMFs. The four IMFs shown have spectral power peaks at about 10 Hz, 18 Hz, 29 Hz, and 40 Hz, respectively.

**Fig. 3.**
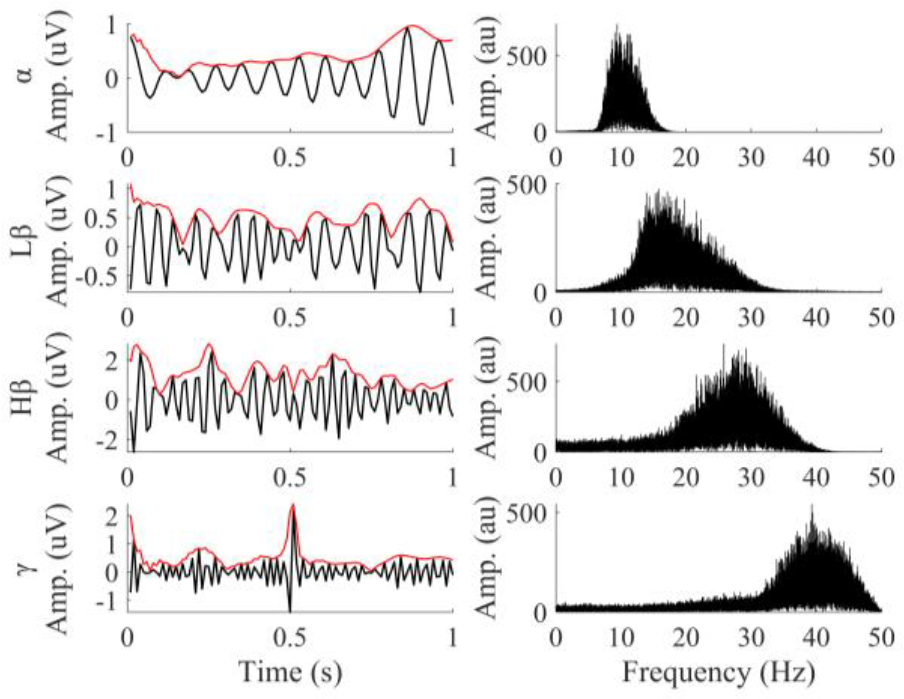
Time series and spectral power of four IMFs extracted from the STN LFP using Masking EMD in Subject 4. The extracted IMFs have frequency spectra corresponding to four canonical LFP frequency bands. The solid red line is the Hilbert envelopes of IMFs.

### 3.3 Assigning HMM States to Canonical Frequency Bands

By assigning HMM states to the canonical frequency bands, we were able to compare states across subjects 35. The upper panel of Fig. 4 shows 12 states that were decoded from HMM analysis in a single patient (Subject 4) during the stepping task. Following the correlation of the state time courses with IMFs, we were able to narrow down the selected states to those that corresponded most to each canonical frequency band. The spectra of states corresponding to each canonical frequency band are shown in the lower panel of Fig. 4.

**Fig. 4.**
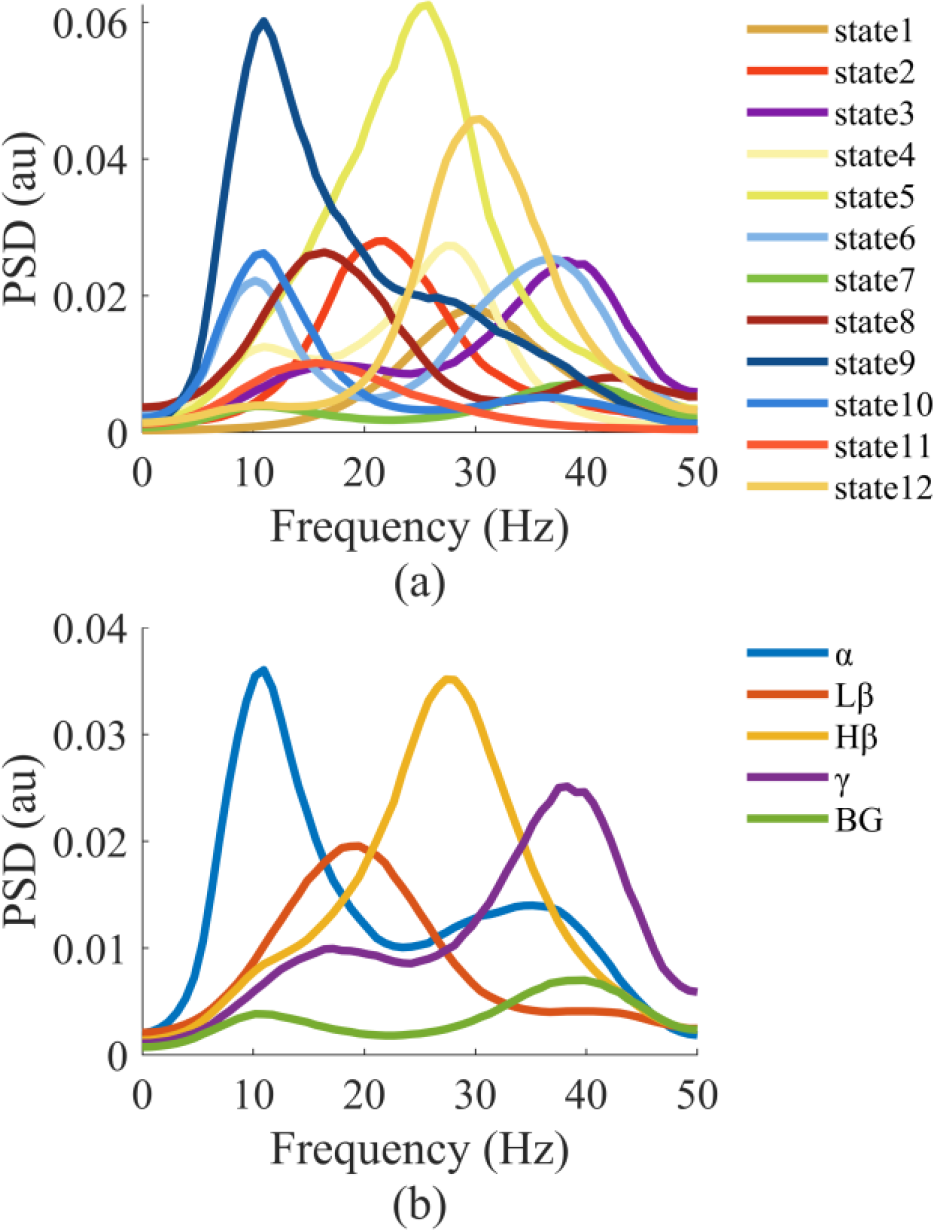
Assigning HMM states to canonical LFP frequency bands. Exemplar data are shown from Subject 4. (a) Upper panel: Spectra of HMM states decoded from the preprocessed LFP are shown. (b) Lower panel: Spectra of the HMM states corresponding to each of the five canonical frequency bands are shown. In (a), the states classified to the same frequency band are averaged to obtain (b) (Lβ: low β, Hβ: high β, BG: background activities).

### 3.4 Temporal Features Dynamics of LFP in Different Bands during Stepping

To probe the influence of different sound conditions on the temporal dynamics of frequency band-specific HMM states, we compared two different time domain features (fractional occupancy and state lifetimes).

Fig. 5 reveals that auditory cues resulted in differential effects on fractional occupancy within the α and Lβ bands. In the α band, fractional occupancy increased in the on-sound and post-sound conditions compared to the pre-sound condition (on vs pre: *p* = 0.0234; post vs pre: *p* = 0.0234). In the Lβ band, fractional occupancy was significantly reduced in the on-sound condition compared to the pre-sound condition (*p* = 0.0234).

**Fig. 5.**
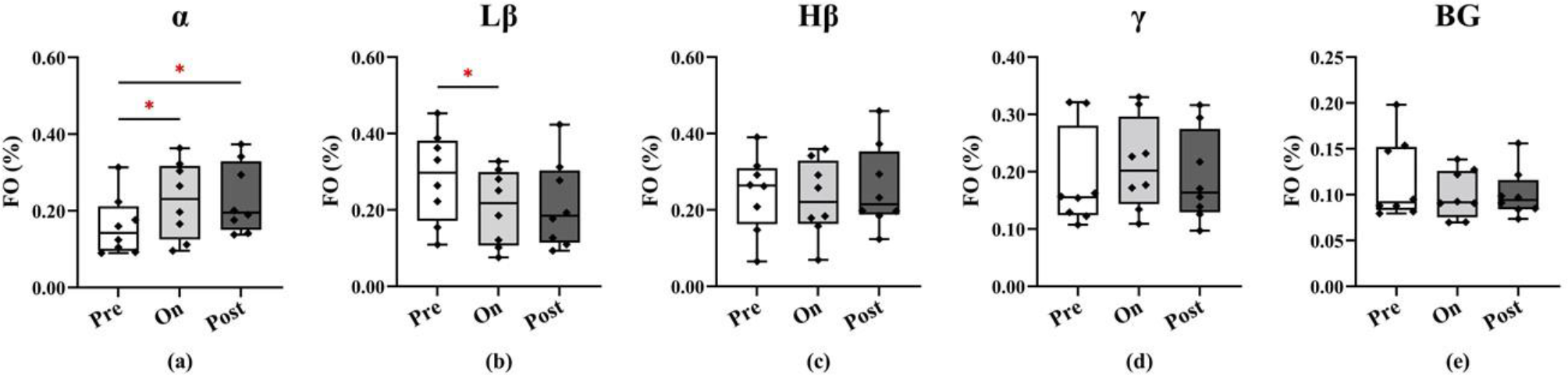
Comparisons of fractional occupancy among the three sound conditions (pre, on, and post sound cues) in the different frequency bands (α, Lβ, Hβ, γ, BG from left to right panels). Each dot represents one subject. *p<0.05, **p<0.01, ***p<0.001.

Effects on state lifetimes are shown in Fig. 6. For the α band, state lifetimes were significantly higher in the on-sound condition compared to the pre-sound condition (on vs pre: *p* = 0.0234).

**Fig. 6.**
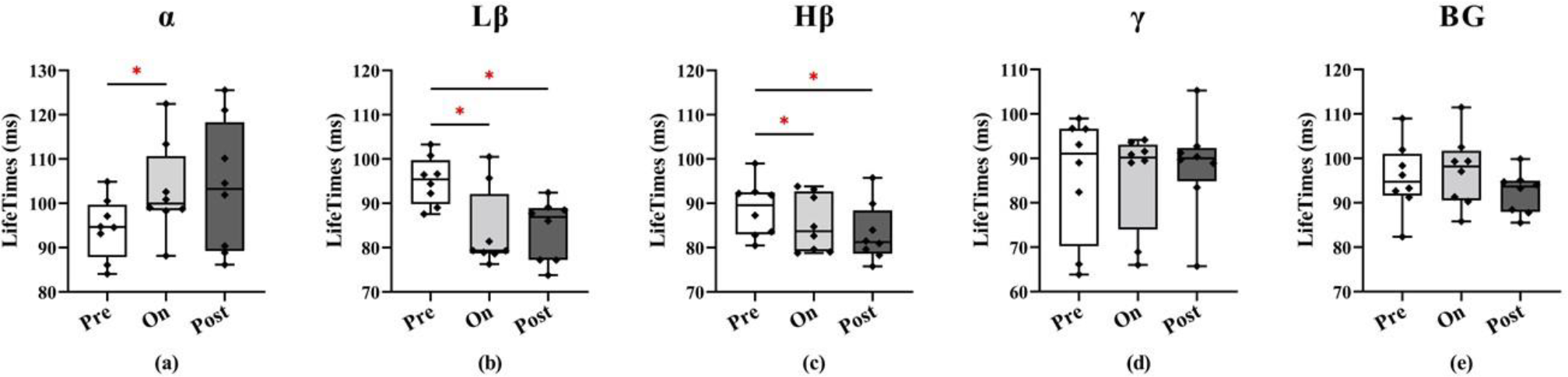
Comparisons of state lifetimes among the three sound conditions (pre, on, and post sound cues) in the different frequency bands (α, Lβ, Hβ, γ, BG from left to right panels). Each dot represents one subject. *p<0.05, **p<0.01, ***p<0.001.

Lβ band state lifetimes were shorter on-sound and post-sound compared to pre-sound (on vs pre: *p* = 0.0352, post vs pre: *p* = 0.0234).

Similar effects were also observed for the Hβ band (on vs pre: *p* = 0.0234, post vs on: *p* = 0.0234). No significant differences were observed in other bands.

### 3.5 The Effect of the Auditory Cue on Transition Probabilities

Next, we examined how transitions between states were influenced during auditory cueing. We explored differences in the transition probabilities between states for all pairwise comparisons of the three different auditory conditions.

Fig. 7 shows the comparison of transition probability differences between the on-sound and pre-sound conditions for the different frequency bands. In the on-sound condition, there was a significant increase in the probability of transitioning from the α band to BG (*p* < 0.001). Additionally, decreased transition probabilities were found for the following state transitions: 1) Lβ to γ (*p* = 0.0122), 2) BG to α (*p* < 0.001).

**Fig. 7.**
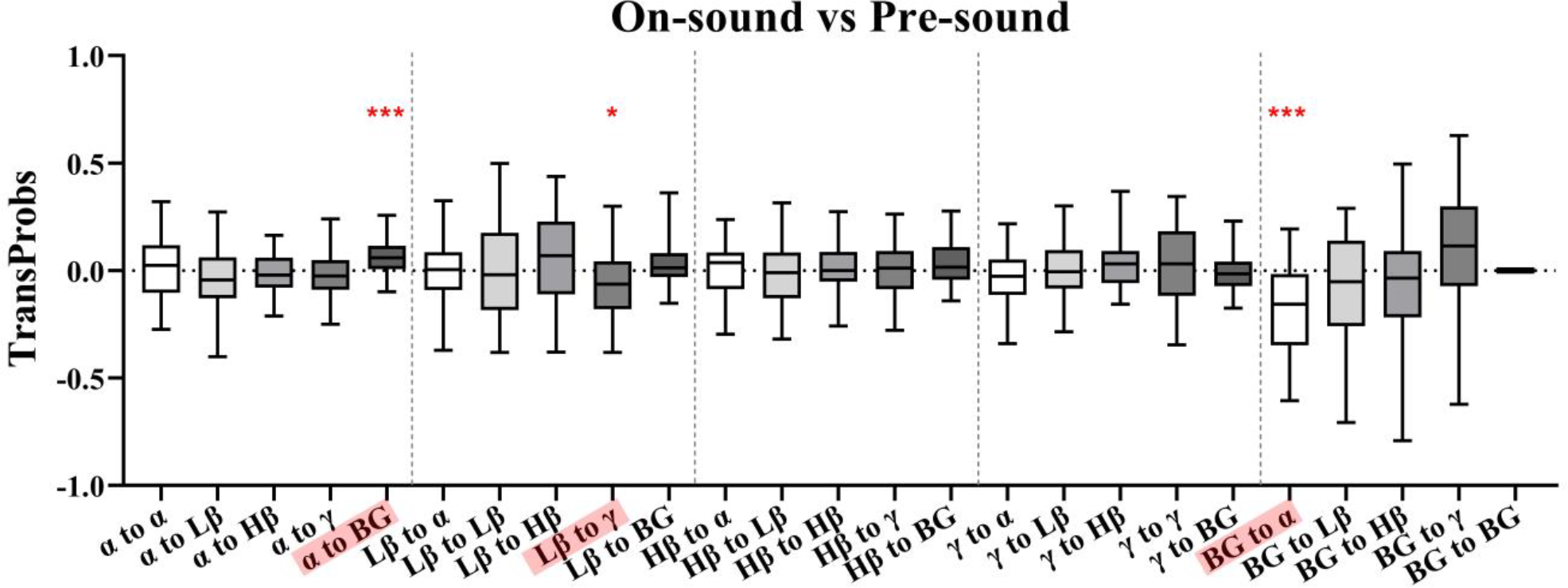
Comparisons of transition probability of the on-sound referenced to the pre-sound conditions in different frequency band pairs. *p<0.05, **p<0.01, ***p<0.001.

Similarly, Fig. 8 shows the comparison of transition probability differences between the post-sound and pre-sound conditions. We only observed a significant increase for the α to BG transitions (α to BG: *p* = 0.0173).

**Fig. 8.**
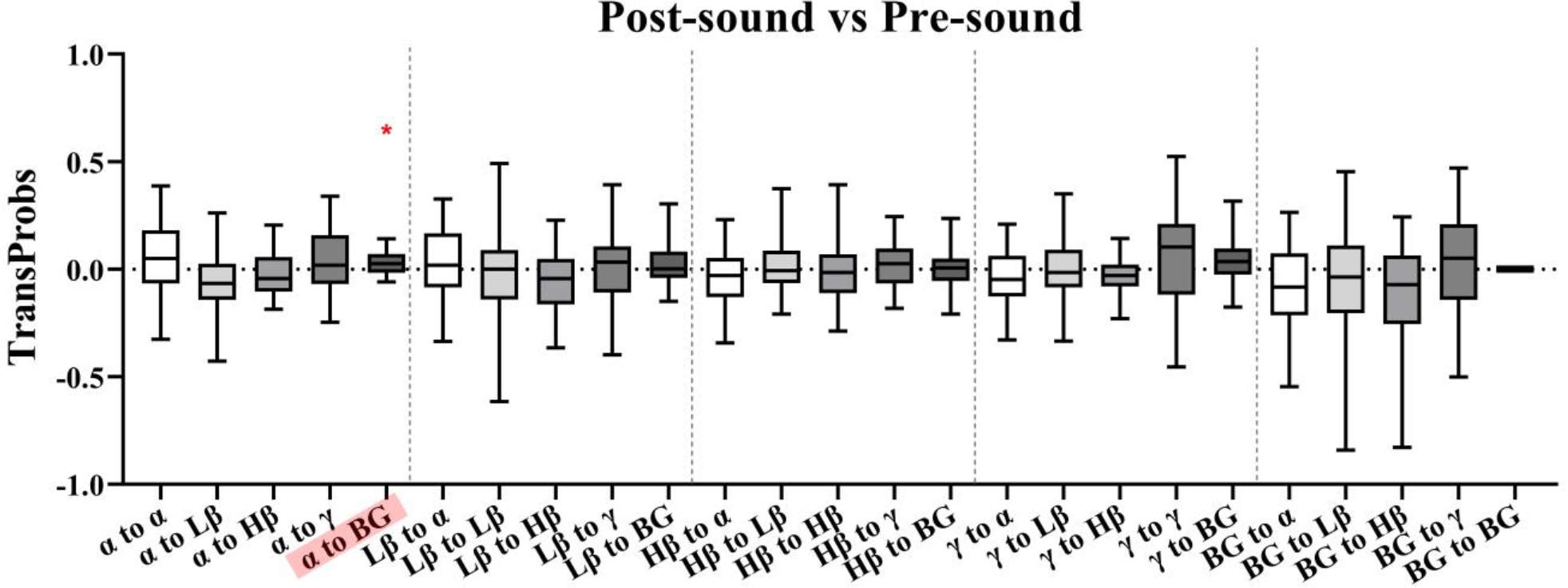
Comparisons of transition probability of the post-sound referenced to the pre-sound conditions in different frequency band pairs. *p<0.05, **p<0.01, ***p<0.001.

Fig. 9 is similar to Fig. 7 and Fig. 8 but compares transition probabilities for the post-sound and on-sound conditions. For the α band, there was a significant decrease in the probability of transitioning to the BG (*p* = 0.0076). For Lβ frequencies, transitions to Hβ were less likely (*p* = 0.0038), whilst transitions to γ were more likely (*p* = 0.0310). Additionally, γ band transitions to Hβ (*p* = 0.0155) were less likely. Finally, we also observed an increase in the probability of transitioning from the BG state to α (*p* < 0.001).

**Fig. 9.**
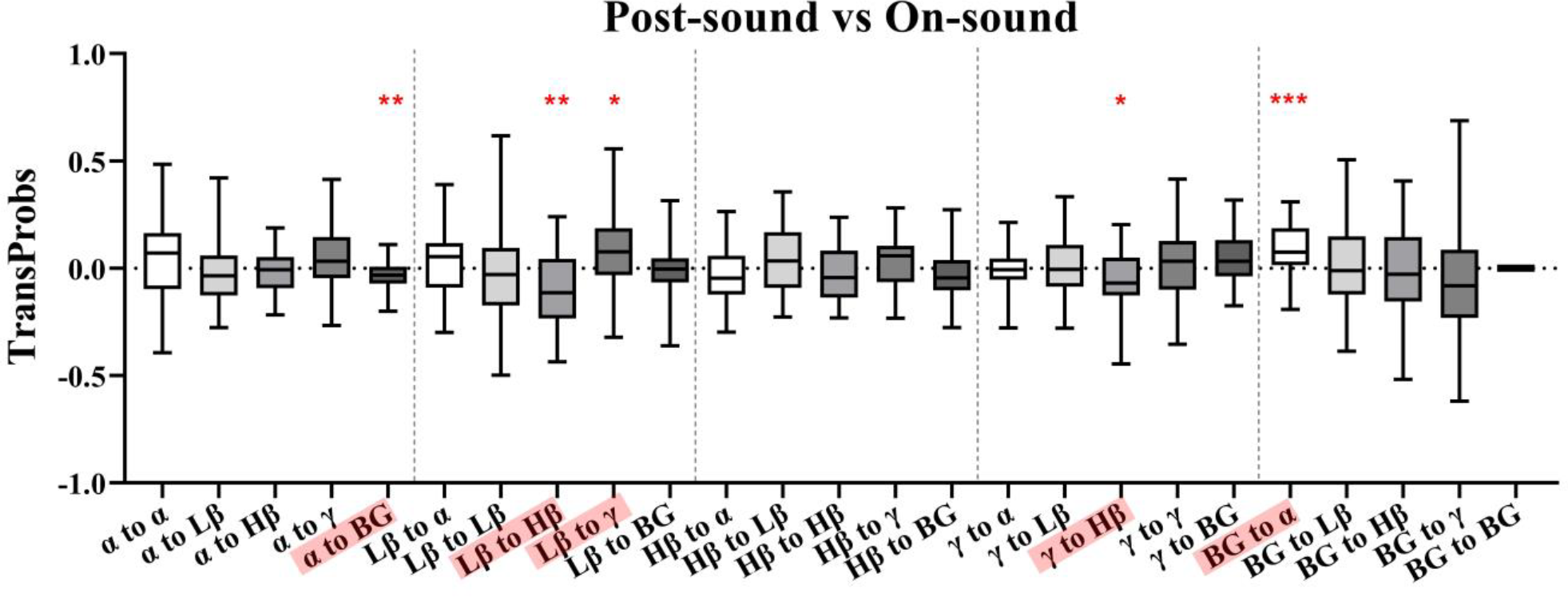
Comparisons of transition probability of the post-sound referenced to the on-sound condition in different frequency band pairs. *p<0.05, **p<0.01, ***p<0.001.

## 4. Discussion

In this study, we develop a novel decoding approach - based on the combination of MEMD and the TDE-HMM for discovering how auditory cues modulate STN activity at short timescales during stepping in patients with PD. We show that auditory cues can influence the properties of states corresponding to activity within multiple frequency bands. Our findings shed light on the mechanisms through which auditory cues improve gait and suggest biomarkers for gait improvement which could be targeted using aDBS strategies. We will first discuss some of the technical nuances of our approach, before focusing on physiological insights.

### 4.1 Performances of MEMD-based TDE-HMM in Decoding STN-LFP states

MEMD is an improved time-frequency analysis method for nonlinear and non-stationary oscillations that is based on standard EMD. In addition to minimizing mode mixing^38^ and mode splitting^37^, MEMD enables the resultant IMFs to more accurately reflect activity within classical EEG bands of interest (i.e., α band, β band, etc. See Fig. 3).

By correlating HMM-identified state time courses with the Hilbert envelope of each IMF, we were able to reliably identify states that captured activity within physiological bands of interest across subjects.

One important limitation of the HMM however is the assumption that different states are mutually exclusive^27,45^. The model selected the state with the highest occurrence probability from multiple states inferred at each time point to form STCs. Brain dynamics are complex, and it may be an oversimplification to assume that multiple states cannot be simultaneously active.

### 4.2 Effect of Auditory Cues and the Roles of Brain States in Different Frequency Bands

Four frequency bands including α, Lβ, Hβ, and γ were considered in this study. We observed that the FO and LTs of β states, particularly Lβ states, were reduced following the occurrence of an auditory cue. Interestingly, auditory cues had a prolonged effect on Lβ and Hβ state LTs, such that they were reduced after the cueing itself (Fig. 5(b) and Fig. 6(b)).

Our findings suggest that auditory cueing may promote gait partly through the suppression of exaggerated STN β band activity. This is perhaps not surprising given that exaggerated β band activity has been closely associated with motoric impairments including gait freezing^10,26^,46-^49^49].

In contrast to the β band, α band states displayed enhanced FO and LTs during and after auditory cues. Although the precise role of alpha band activity is unclear, recent reports suggest that synchronizing activity within this frequency range may produce prokinetic effects^50^. Another equally plausible possibility is that α band synchronisation may support attentional and executive processing involved in gait and motor preparation^51^. γ band activities are closely linked to movement, and deficiency of fast γ bursts was reported to contribute to bradykinesia^52^. Interestingly we observed no auditory cue related changes in FO and LTs corresponding to γ band activity, only slight changes in γ band were observed during state transitions, which may be due to the relatively constrained nature (stepping in pace while sitting) of the task.

Finally, we observed that auditory cues also modulated the transition probabilities of individual states, including the α, Lβ, γ, and background activity bands.

The reduced transition probability from background activity to alpha states indicates that alpha states were more sustained, or potentially stable as also shown in the life time analysis. However, we also observed an increased probability of alpha to background state transitions, which was most pronounced when the sound was present and persisted – although weaker – during post-sound stepping. An increased likelihood of transitioning to BG activity should go in hand with a reduced likelihood of transitioning from alpha to beta and gamma states, however the differences were only moderate and not significant, i.e., not sufficiently consistent.

Interestingly, we saw a reduced probability of state transitions from low beta to gamma in the presence of the sound, however considering trends of decrease in the probability of transition from alpha and BG to low beta, likely a reflection of the relative reduction of low-beta states as a whole. Perhaps it is exactly this transition that might make patients susceptible to freezing of gait in the absence of auditory cues. These trends of differences were maintained in the post-sound period.

Previous analysis of the same data has shown that high beta activity was particularly strongly modulated by the left-right alternating stepping cycle, and that bursts around 30 Hz were more likely during the contralateral stance period, which was further enhanced with auditory cueing [9]. Splitting the high beta band further into 20-25 and 26-30 Hz might have resulted in more pronounced changes in the higher Hβ band. However, correcting for an even larger number of multiple comparisons would have reduced our chance of identifying small effects.

The observed changes in state transition probabilities may be the mechanism that underlies the observed changes in FO and state LTs for the α and β frequency bands. Future studies with larger sample sizes could enable more detailed examinations of the composition of what we defined as “background activity state” or a more fine-grained analysis of high beta states.

### 4.3 Limitations

Several limitations should be highlighted for interpreting the results of this study. Firstly, LFP recordings could potentially have been affected by microlesions caused by the electrode insertion procedure. Secondly, LFPs were recorded while patients were seated, performed stepping-in-place movements, and did not experience freezing of gait. Somewhat reassuringly, previous analysis of a subset of data from the same subjects included in this study has revealed very similar STN spectral modulations during stepping while standing, and free walking^9^. This serves to highlight the potential applicability of our findings to cued free walking. Finally, although the sample size of this study was limited to only eight patients, we were able to reveal significant cue related modulations of both α and β activity.

## 5. Conclusions

We have combined MEMD and the TDE-HMM to decode dynamic brain states during cued stepping in patients with PD. Our results indicate that auditory cues, which are known to improve gait performance, exert differential effects on STN LFP activity within the α and β bands. Specifically, auditory cues led to the enhancement of α band activity, whilst reducing the probability of β synchronization states. Our findings reveal a potential neurophysiological mechanism for the effects of auditory cues and could help inform therapeutic aDBS strategies that could be tailored to intermittently enhance α band activity while suppressing β.

## Data availability

The datasets used for the current study are available from the corresponding author on reasonable request (https://data.mrc.ox.ac.uk/data-set/).

## Acknowledgments

This work is supported by the National Natural Science Foundation of China (62001026, 62171028), the Open Project of Key Laboratory of Medical Electronics and Digital Health of Zhejiang Province (MEDH202204, MEDC202303), and the BIT High-level Fellow Research Fund Program (3050012222022). H.T. is supported by the Medical Research Council UK (MC_UU_00003/2, MR/V00655X/1, MR/P012272/1), the National Institute for Health Research (NIHR) Oxford Biomedical Research Centre (BRC) and the Rosetrees Trust, UK. A.O. is supported by an MRC Clinician Scientist Fellowship (MR/W024810/1).

## Author contributions

C.-H.Y. and W.S. contributed to the conception and methodological design of the study; J.J.F., A.L.G., P.F. and H.T. contributed to the acquisition of the data; C.-H.Y. and Y.X. contributed to the code implement; C.-H.Y., Y.X., P.F., H.T. and A.O. contributed to the analysis of the data and the interpretation of results; C.-H.Y. and Y.X. drafted the manuscript; C.-H.Y., W.S., J.J.F., A.L.G., P.F., H.T. and A.O. revised the manuscript, and C.-H.Y., W.S., H.T. and A.O. obtained funding. All authors read and approved the final manuscript.

